# Disease investigation in a New Zealand high-health pig herd reveals a divergent porcine adenovirus

**DOI:** 10.64898/2026.09.14.751650

**Authors:** Stephanie J Waller, David J Pulford, E Bruce M Welch, Bernard L Vaatstra, John P O’Connell, Kimiya Byrne, Hayley Hunt, Harry S Taylor, Jemma L Geoghegan

## Abstract

Porcine adenoviruses are common in domestic pigs and are generally regarded as low-pathogenicity viruses associated with mild or subclinical infections. However, their diversity, host associations and disease potential remain poorly understood. Here, we investigated a disease outbreak in an immunologically naïve 320-sow high-health hysterectomy-derived pig herd in New Zealand. The outbreak was characterised by anorexia, lethargy, diffuse cutaneous erythema, death in neonatal piglets, and an increased occurrence of mummified fetuses. Histopathological examination of a piglet revealed widespread intranuclear inclusion bodies within endothelial cells, indicative of systemic viral infection. Total RNA sequencing was performed on tissues from an affected piglet and mummified fetuses. Metatranscriptomic analysis identified an abundant adenovirus in multiple piglet and aborted fetus tissues, and *de novo* assembly recovered a 27,685-nucleotide genome. Phylogenetic analysis placed the virus within the genus *Mastadenovirus* but it formed a divergent lineage distinct from all previously characterised porcine adenoviruses. Despite sequence divergence, the genome retained the characteristic organisation of mastadenoviruses. Aside from a single detection of porcine kobuvirus identified at low abundance in the small intestine and colon from the piglet, no other viral pathogens were identified. The high viral abundance and systemic tissue distribution observed in both the piglet and fetus, together with the presence of widespread adenoviral inclusion bodies in the piglet, support an association between this novel adenovirus and the disease outbreak. These findings expand the diversity of porcine adenoviruses and suggest that previously unrecognised adenovirus lineages may contribute to reproductive and neonatal disease when introduced into immunologically naïve high-health herds.

**Importance:** Porcine adenoviruses are widespread in pigs, but their ability to cause disease is not well understood. We investigated an outbreak of severe disease and reproductive losses in a high-health herd of domestic pigs in New Zealand and identified a novel porcine adenovirus. The virus was found at high abundance across multiple tissues from an affected piglet and fetuses, that showed widespread tissue damage consistent with adenovirus infection. The virus was also genetically distinct from previously described porcine adenoviruses, revealing an unrecognised lineage within the group. These findings expand our understanding of adenovirus diversity in domestic pigs and highlight that previously undetected viruses may cause serious disease when introduced into immunologically naïve, high-health herds. The lack of prior exposure to viruses may have contributed to the severity of disease observed in this outbreak. Identifying such viruses is important for understanding emerging diseases and protecting high-health pig populations.

## Introduction

Adenoviruses, belonging to the family *Adenoviridae*, are non-enveloped viruses with linear double-stranded DNA genomes ranging from 25-48 kb in length^1^. The family is divided into six genera: *Mastadenovirus* (mammals), *Aviadenovirus* (birds), *Atadenovirus* (reptiles, birds, ruminants and marsupials), *Siadenovirus* (birds, frogs and tortoises), *Ichtadenovirus* (fish) and *Testadenovirus* (terrapins)^1^. Human adenovirus infections are typically mild or asymptomatic, but can cause respiratory^2^, ocular^3^, gastrointestinal^4^, hepatic^5^ and urinary tract disease^6^, particularly in immunocompromised individuals. In animals, adenoviruses are widespread and often associated with subclinical infections, although some can cause severe disease, including infectious canine hepatitis in dogs^7^, hepatitis and gizzard erosion in chickens^8^, egg drop syndrome in poultry^9^, haemorrhagic enteritis in turkeys^10^, fatal enteric disease outbreaks in young cattle^11^ and fatal disease outbreaks in deer^12^.

Porcine adenoviruses belong to the genus *Mastadenovirus* and are widespread in domestic pig populations but are generally considered to be of low pathogenicity^13^. Although five porcine adenovirus serotypes have been recognised, genomic data remain limited, restricting our understanding of their evolutionary diversity, host associations and disease potential^13,14^. Porcine adenovirus was first isolated in 1964 from a pig with diarrhoea in England^15^ and has since been reported worldwide^13^. In New Zealand, prior to the advent of molecular diagnostics, inclusion bodies attributed to adenovirus have been an incidental finding on histological examination of piglets^16^. Subsequently Porcine adenovirus 3 (PAdV-3) and PAdV-5 have been detected in pig faecal samples during evaluation of a multiplex RT-qPCR assay, while PAdV-5 was also detected in river water and shellfish as a marker of porcine faecal contamination^17^. However, to date, no porcine adenovirus-associated disease outbreaks have been reported in New Zealand.

Most porcine adenovirus infections are associated with mild, self-limiting diarrhoea^14^, although the virus has also been detected in clinically healthy pigs without apparent signs of disease^13,18^. Transmission is believed to occur primarily via the faecal-oral route^13^. Porcine adenoviruses have occasionally been identified in pigs with respiratory disease^19,20^, and in one case of a pig with encephalitis^21^, although their role in the pathogenesis remains uncertain. Reports of systemic porcine adenovirus infection are rare. However, a naturally occurring disseminated adenovirus infection has been described in a nursing pig in North America, characterised clinically by haemorrhagic diathesis of the skin and histologically by widespread viral inclusions within endothelial, interstitial and epithelial cells across multiple tissues, including the skin, kidney, spleen, liver, heart and small and large intestines^22^. Severe systemic disease has also been reported in hysterectomy-derived, colostrum-deprived piglets, which developed skin cyanosis and widespread eosinophilic intranuclear inclusion bodies within endothelial cells of capillaries and small blood vessels throughout the body ^23^. Ultrastructural examination revealed adenovirus-like particles within affected nuclei, suggesting that the virus may have been transmitted transplacentally^23^. Experimental infections in germ-free and pathogen-free pigs have further demonstrated the ability of porcine adenoviruses to disseminate systemically, with lesions observed in multiple organs, including the lungs, kidneys, thyroid and lymphoid tissues^21^.

In April 2025, a disease outbreak occurred in a 320-sow high-health pig herd in New Zealand established from hysterectomy-derived piglets. Such high-health herds are maintained free of a range of endemic pathogens through strict biosecurity and ongoing health surveillance, making the emergence of unexplained disease particularly concerning. Approximately half of the piglets from two litters, aged 5-6 days, developed clinical signs including anorexia, lethargy, and diffuse reddening of the skin (Figure 1), raising suspicion of a septicaemic process. Affected piglets showed no evidence of diarrhoea. Concurrently, the proportion of mummified fetuses farrowed each week increased markedly. Here, we describe the diagnostic investigation of this outbreak and the identification of a novel porcine adenovirus associated with systemic disease.

**Figure 1.**
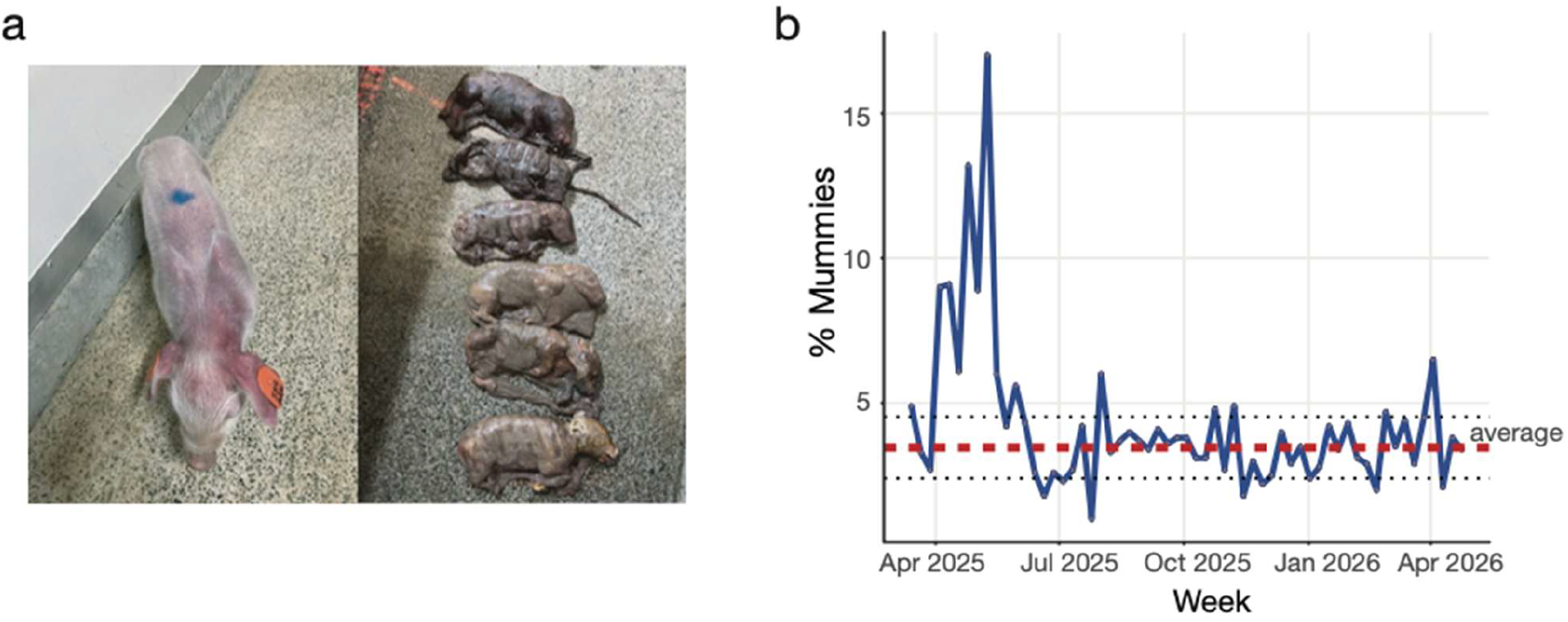
**(a)** Tissues from an untreated piglet presenting with a suspected septicaemic condition (left) and from mummified fetuses (right) were collected during an investigation of a disease outbreak on a New Zealand pig farm during April 2025. **(b)** Percentage of mummified fetuses per week at the affected farm. The red dashed line indicates the mean percentage of mummies from July 2025 onwards, and the black dotted line represents the standard deviation from July 2025 onwards.

## Methods

### Case History

A 320-sow (*Sus scrofa domesticus*) high-health New Zealand herd that was assembled in 2020 from hysterectomy-derived powdered bovine colostrum and milk reared piglets, with a view to excluding significant bacterial pathogens of swine including *Streptococcus suis*, experienced a disease outbreak in April 2025.

The herd typically farrowed approximately 14 litters per week. Over a one-week period in mid-April 2025, up to half of the piglets from two litters of 19 and 21 piglets were affected at 5-6 days of age. Clinical signs included anorexia, lethargy and diffuse reddening of the skin. A septicaemic condition was suspected, and affected piglets were treated albeit unsuccessfully with 30 mg/kg of long-acting amoxycillin (Vetrimoxin LA, Ceva Animal Health (NZ) Ltd.,Auckland, NZ). Only two litters were confirmed to be affected, although there were unverified reports of a third litter from the same farm being affected in the same week.

Coinciding with the outbreak, the proportion of mummified fetuses farrowed per week increased markedly, rising to 9.0% in the week ending 4 April 2025 and peaking at 17% in the week ending 9 May 2025 (Figure 1). This was substantially higher than the average weekly prevalence of 3.5%. The proportion subsequently declined to 6% in the week ending 16 May 2025, returning towards baseline levels (Figure 1). All other production and health parameters at the time of the outbreak were considered normal, including pre-wean mortality and the number of sows that failed to farrow.

### Sample Collection

One untreated piglet (Piglet 1) exhibiting clinical signs of anorexia, lethargy and diffuse reddening of the skin was euthanised and a postmortem examination was performed. Aside from the skin lesions, the only notable postmortem finding was mesocolonic edema. Tissues (lymph node, spleen, kidney, liver, heart, lung, small intestine and colon) were collected in 10% buffered formalin for diagnostic investigation (Figure 1, Supplementary Table 1). Additionally, fresh lung, thymus, liver, kidney, spleen, heart and brain samples from four fetuses (Fetus 1-4) were collected, tissues were homogenised in PBS at 10% w/v using Bioreba extraction bags (Bioreba AG), then centrifuged at 1,000 x g and supernatant was stored at - 80°C (Supplementary Table 1). A further four fetuses (Fetus 5-8) were frozen at -20°C for 8 months before kidney, liver, lung and spleen samples were harvested and stored separately in 800ul of RNA stabilisation solution (DNA/RNA Shield, Zymo Research) at -80°C until RNA was extracted (Supplementary Table 1).

### Histopathology and electron microscopy

The formalin-fixed piglet samples of heart, liver, kidney, spleen, lymph node, lung, small intestine and colon were processed routinely for histological examination at a commercial veterinary diagnostic laboratory. For electron microscopy, 2mm cubes of the spleen and small intestine were removed from the paraffin blocks used to prepare the histological sections. The paraffin was melted by briefly placing the cube in a warm (70°C) oven, then the tissue cubes were processed through four changes of xylene before being moved through decreasing concentrations of ethanol and finally placed in buffer, as described by Graham and Orenstein (2007)^24^. The cubes were post-fixed in osmium tetroxide and embedded into epoxy resin. Thin sections from the resin blocks were stained using lead citrate/uranyl acetate and viewed using an FEI Tecnai G2 Biotwin Transmission Electron Microscope (Hillsboro, OR, USA) at the Manawatū Microscopy and Imaging Centre (Massey University).

### Nucleic acid extraction and PCR screening

Nucleic acids were extracted from 100 µL of tissue supernatant using the MagMAX™ CORE Nucleic Acid Purification Kit (Thermo Fisher Scientific) according to the manufacturer’s instructions and eluted in 90 µL. Nucleic acids were also extracted from formalin-fixed paraffin-embedded (FFPE) tissue scrolls from the piglet using the NucleoSpin® DNA FFPE XS kit (Macherey-Nagel) and eluted in 20 µL. Scrolls from three FFPE tissue blocks were processed: the first contained spleen and lymph node, the second contained heart, lung, liver, and kidney, and the third contained small intestine and colon.

All eluted nucleic acid extracts were screened using an 18S rRNA TaqMan qPCR for assessing nucleic acid quality for PCR. Samples were subsequently tested for herpesviruses, adenoviruses and porcine circoviruses using previously published pan-herpesvirus, pan-adenovirus, porcine adenovirus and porcine circovirus PCR assays. Conventional and nested PCRs were performed using KAPA2G Fast HotStart ReadyMix (Kapa Biosystems), hydrolysis probe PCR assays used the SuperScript® III Platinum® One-Step qRT-PCR System (Thermo Fisher), and SYBR Green assays used SYBR® Green JumpStart Taq ReadyMix (Sigma-Aldrich). Primer sequences and cycling conditions are provided in Supplementary Table 2.

### Total RNA extraction for metagenomic analysis

Excess paraffin wax was removed from FFPE tissue scrolls from piglet 1 before deparaffinisation. Three tissue scrolls were processed: the first contained spleen and lymph node, the second contained heart, lung, liver, and kidney, and the third contained small intestine and colon. Samples were then deparaffinised and digested at 55°C for an hour and total RNA was extracted using the Zymo Quick-RNA FFPE Kit (Zymo Research). Fetal tissues from fetus 5-8 (kidney, liver, lung and spleen; four tissues per piglet) were defrosted before 10mg of tissue were placed in ZR BashingBead Lysis Tubes (0.1 mm and 0.5 mm) (Zymo Research) filled with 750ul of DNA/RNA shield (Zymo Research). Lysis tubes were placed into a mini-beadbeater 24 disruptor (Biospec Products Inc.) and were homogenized for five minutes. Total RNA was extracted using the ZymoBIOMICS MagBead RNA kit (Zymo Research) following the manufacturers protocol. RNA was quantified using a NanoDrop (Thermo Fisher Scientific). Equal volumes of RNA from each of the tissue types were pooled by pig into four pools. In total, RNA from three FFPE tissue blocks from piglet 1 and four pooled fetal tissue samples (one from each of four fetuses 5-8, each comprising kidney, liver, lung, and spleen) were sent for sequencing.

### RNA sequencing

Extracted RNA was subject to total RNA sequencing. Libraries were prepared using the Illumina Stranded Total RNA Prep with Ribo-Zero Plus (Illumina) and 16 cycles of PCR. Paired-end 150 bp sequencing of the RNA libraries was performed on the Illumina NovaSeqX platform.

### Virome assembly and virus identification

Initial read quality was assessed using FastQC^25^. Sequencing reads were quality-trimmed using Trimmomatic v0.38^26^. Adaptor sequences were removed, low-quality bases with quality scores of less than 3 were trimmed from the 5′ and 3′ ends, and sliding window trimming (4 bases, average quality <5) was applied. Reads shorter than 25 nucleotides after trimming were discarded and only paired reads retained after trimming were used for downstream analyses. Post-trimming read quality was reassessed using FastQC to confirm improvement in sequence quality^25^. Following quality control, reads were *de novo* assembled into contigs using MEGAHIT v1.2.9^27^ using default parameters. Contigs were then compared against the NCBI non-redundant protein (nr) databases using DIAMOND v2.1.9^28^ to identify putative viral sequences based on sequence similarity to known viruses.

### Estimating viral transcript abundance estimations

Virus abundance was estimated by mapping reads back to the assembled contigs using Bowtie2 v2.4.4^29^ and SAMtools v1.9^30^ using the local alignment mode (--local) with default parameters. Total viral abundance estimates for *Porcine adenovirus* was compiled across all libraries. Estimated abundances per million reads were standardized to the number of paired reads per library.

### Virus phylogenetic analysis

Translated conserved adenovirus DNA polymerase protein sequences were aligned with representative protein sequences from the same virus family obtained from NCBI RefSeq as well as the closest BLASTp hits using MAFFT v7.490^31,32^ using the auto algorithm and default settings. Ambiguously aligned regions were removed using trimAL v1.2rev59^33^ with the gap threshold flag set to 0.9. Phylogenetic trees for each viral species/family/order were then estimated using the maximum likelihood method in IQ-TREE v1.6.12^34^, employing the LG amino acid substitution model with 1000 ultra-fast bootstrapping replicates. The resulting phylogenetic tree was annotated using Figtree v1.4.4^35^.

### Temporal analysis of mummified fetuses

Weekly production records were used to investigate temporal changes in the proportion of mummified fetuses. Temporal trends in the percentage of mummified fetuses were visualised using line plots generated with the ggplot2 package^36^ in R version 4.6.0. To assess deviations from baseline production levels, the mean percentage of mummified fetuses and the corresponding standard deviation (SD) from were calculated from July 2025 onwards. Horizontal reference lines representing the mean and mean ± 1 SD were overlaid on the time-series plot to facilitate identification of periods with elevated levels of mummification relative to historical farm performance.

### Viral abundance analysis

Viral abundance was quantified as reads per million sequenced reads (RPM) for each library. Viral abundance was visualised as grouped bar plots using ggplot2^36^, with libraries displayed on the x-axis and RPM values on the y-axis. Because viral abundances spanned several orders of magnitude and included zero values, one was added to all RPM values prior to log10 transformation (RPM + 1). This transformation enabled simultaneous visualisation of both highly abundant and low-abundance viral taxa while retaining libraries in which a virus was not detected.

## Results

Histopathological examination of tissues from the infected piglet revealed widespread intranuclear inclusion bodies consistent with systemic infection. Inclusion bodies were identified predominantly within vascular endothelial cells, macrophages, and stromal cells of the small intestine, colon, lymph nodes, liver, kidney, and spleen (Figure 2a-c). These inclusions were deeply eosinophilic to amphophilic and associated with margination of nuclear chromatin. Occasional inclusion bodies were also observed within hepatocytes and renal tubular epithelial cells. The distribution of inclusion bodies across multiple organs, particularly within endothelial cells, indicated systemic viral dissemination. On electron microscopy, inclusions within stromal cells of the small intestine were composed of clusters of round to hexagonal, electron-dense viral particles measuring 40-70 nm in diameter (Figure 2d). Nuclei containing these particles also exhibited infolding of the nuclear membrane and margination of chromatin.

**Figure 2.**
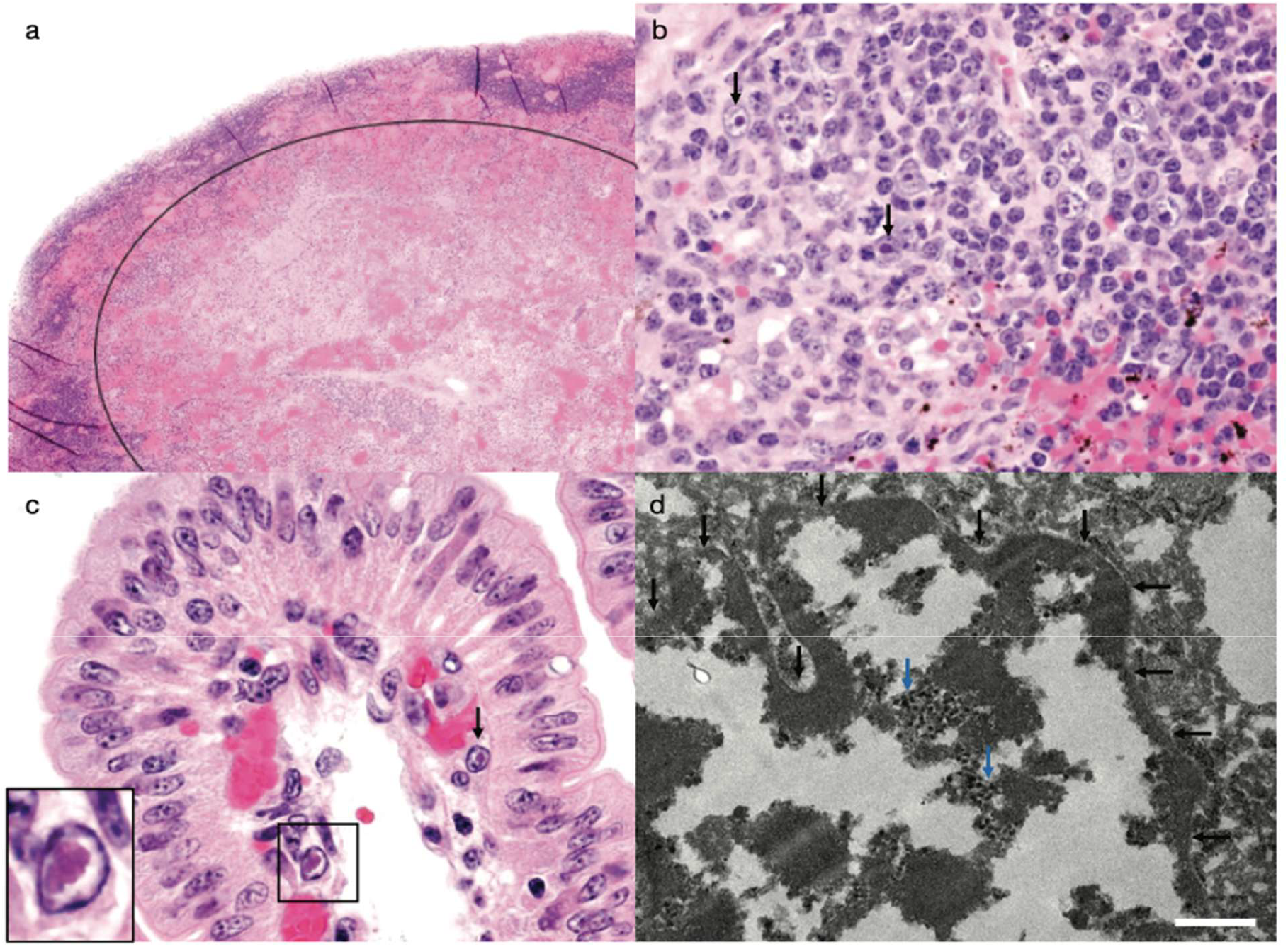
Histopathology and electron microscopy of piglet tissues. (a) Lymph node, with a large area of haemorrhage and necrosis inside the black circle, HE 20x; (b) Higher power view of the lymph node, with amphophilic intranuclear inclusions in macrophages (black arrows), HE 400x; (c) Small intestinal mucosa, with viral inclusions in nuclei of endothelial cells in the lamina propria (black arrow and inset). The inclusions are eosinophilic to amphophilic and marginate the chromatin, HE 400x; (d) Electron microscopy of a stromal cell in the small intestine. The nuclear membrane is indicated by black arrows and is folded. Clusters of round to hexagonal viral particles are present within the nucleus (blue arrows). Scale bar = 500nm.

In addition to the inclusions, on histology the small intestine also exhibited mild villous blunting with multifocal aggregates of basophilic granular material (mineral) within the villi. In the colon, multifocal degeneration and necrosis of the luminal epithelium were evident, characterised by epithelial vacuolation and karyorrhexis. Adherent clusters of bacterial rods and filamentous organisms were present along the epithelial brush border. The lymph node was disrupted by focally extensive areas of necrosis and haemorrhage, while rare renal tubules contained cellular debris. Collectively, these findings were consistent with a systemic viral infection accompanied by enteric and lymphoid pathology.

Fetal tissues were screened by PCR for a panel of DNA and RNA viruses associated with porcine reproductive failure and mummification, including porcine circovirus, porcine reproductive and respiratory syndrome virus, porcine cytomegalovirus, porcine parvovirus type 1 and porcine teschovirus. Porcine circovirus DNA was detected in low quantities by qPCR in fetus 4, with PCR results consistent with infection by porcine circovirus type 1 (PCV1) and type 2 (PCV2), both of which are endemic in New Zealand^37^. Otherwise, all samples were negative for these viral targets (Supplementary Table 3 and Supplementary Figure 1).

A pan-adenovirus nested PCR was applied to all fetal tissues. Amplification products consistent with adenoviral targets were detected in multiple tissues from fetuses 1-3, whereas no amplification was observed in fetus 4 (Supplementary Table 4, Supplementary Figure 1b). Sequence data from adenovirus amplicons was used to design confirmatory porcine adenovirus–specific PCR assays, (PAdV-short and long) and they consistently amplified samples from fetus 1-3 and not fetus 4 to confirm adenovirus positive targets (Supplementary Figure 1c).

Total RNA sequencing of the piglet tissues and four pooled fetal tissue samples (fetuses 5-8) identified a novel porcine adenovirus, provisionally named Porcine adenovirus 6 (PAdV-6) (PZ721986). PAdV-6 contigs were detected in all pooled tissue samples from the piglet at high abundance, ranging from 14,518-26,245 reads per million (RPM) (Figure 3). The virus was also detected in two of four fetal tissue samples, although at lower abundance (22-547 RPM). Phylogenetic analysis of the conserved DNA polymerase gene (1,039 amino acids in length; spanning nucleotide positions 4,020-7,136 of the genome) revealed that this virus fell within the *Mastadenovirus* genus and was novel as highlighted by the divergent lineage (Figure 4). The closest amino acid matches were the DNA polymerases of Polar bear adenovirus 1 (AWY10570.1), previously identified in a 4-month-old polar bear (*Ursus maritimus*) from Germany^38^ and Bat mastadenovirus (WXG22592.1), identified in intermediate roundleaf bats (*Hipposideros larvatus*) from China^39^, both sharing 63% amino acid sequence identity with PAdV-6. The absence of close phylogenetic clustering with known porcine adenoviruses was unexpected. Within the conserved DNA polymerase region, the virus shared only 49% and 60% amino acid identity with PadV-3 and PadV-5, respectively, suggesting that the diversity of adenoviruses circulating in pigs and other mammals may be underrepresented in currently available genomic data. Comparison of the translated open reading frames (ORFs) further demonstrated that PAdV-6 is highly divergent across the genome, with variable amino acid identity to previously described mammalian mastadenoviruses (Supplementary Table 5).

**Figure 3.**
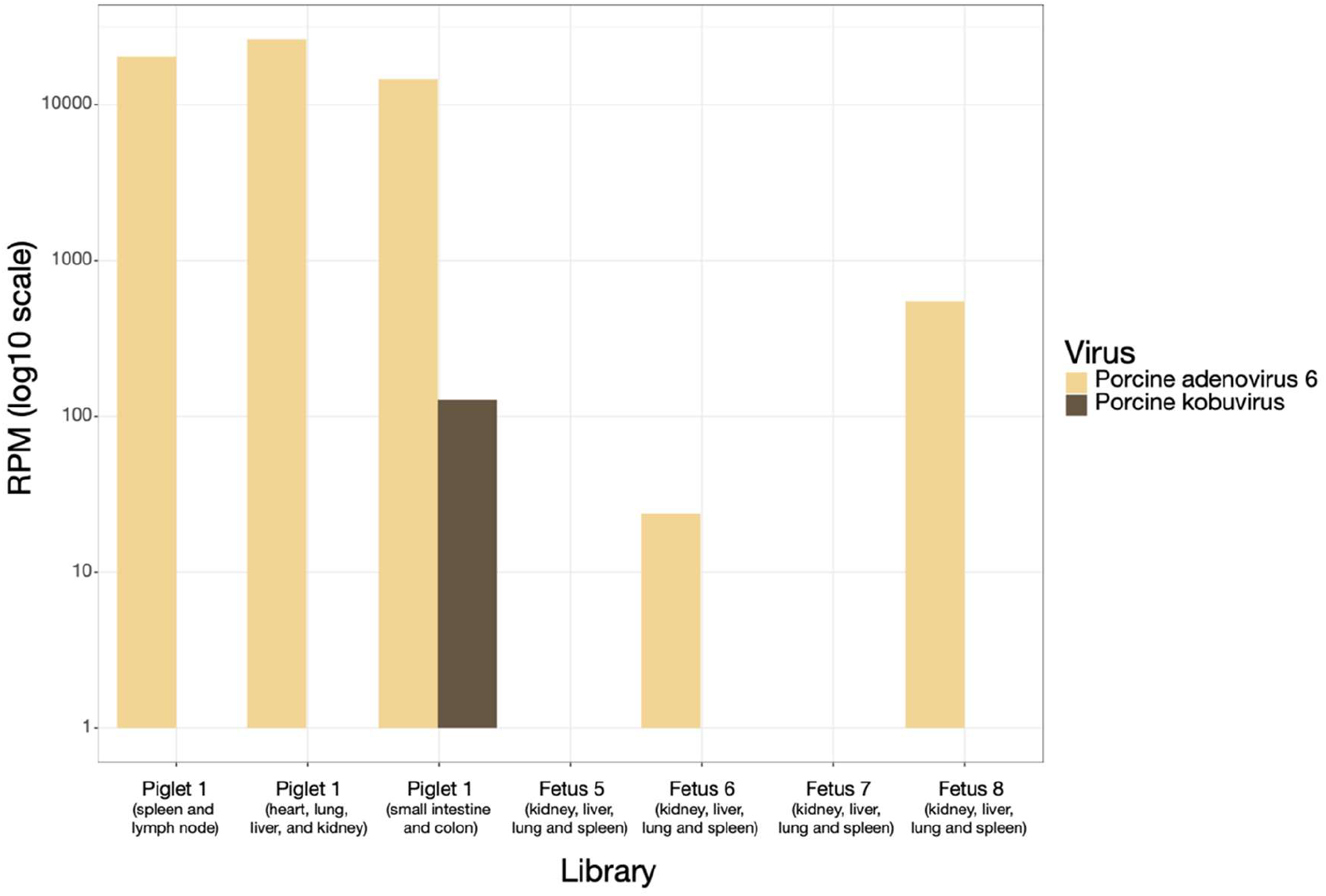
Relative abundance of viral reads assigned to Porcine adenovirus 6 and Porcine kobuvirus across the seven porcine tissue libraries. Viral abundance is expressed as reads per million (RPM) and displayed on a log_10_-transformed scale following the addition of one read (RPM + 1) to accommodate zero values. Bars represent the abundance of each virus detected in individual libraries, including samples from the piglet and fetus samples.

**Figure 4.**
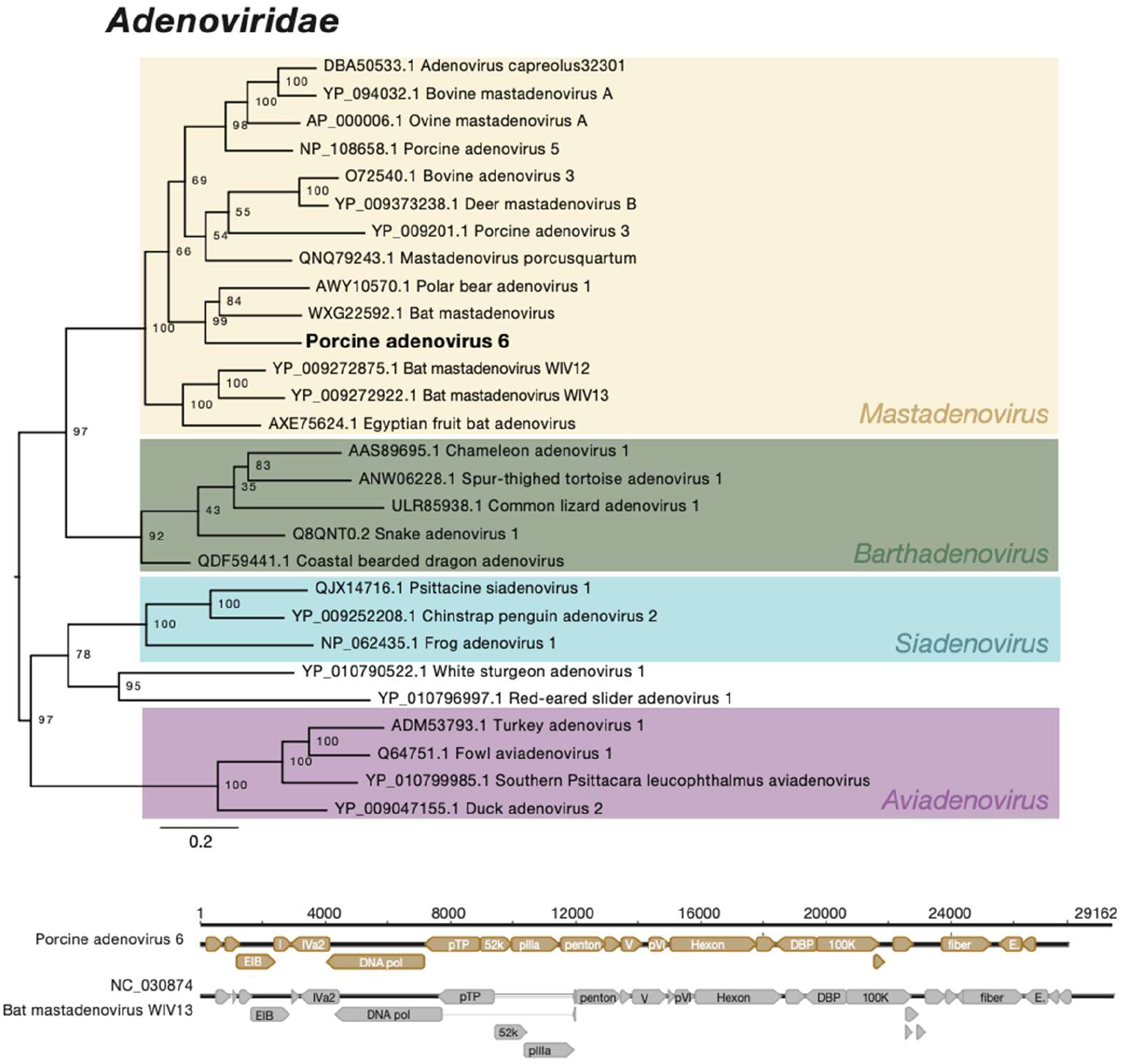
Phylogenetic tree and genome structure of the novel porcine adenovirus. A maximum likelihood phylogenetic tree was constructed from representative adenovirus DNA polymerase transcripts. Branch lengths are proportional to the number of amino acid substitutions per site, and the tree is midpoint rooted. Ultrafast bootstrap values are noted and known genera are coloured. Beneath the phylogenetic tree is the genome structure of the novel pig adenovirus compared with the genome structure of Bat mastadenovirus WIV13 (NC_030874.1).

At the nucleotide level, the closest matches to the 27,685 nt genome were Bat mastadenovirus WIV12 (NC_030860.1) and Bat mastadenovirus WIV13 (NC_030874.1) previously discovered in common bent-wing bats (*Miniopterus schreibersi*) from China, sharing 76% and 73% nucleotide identity, respectively^40^. However, these matches covered only 11% and 18% of the genome, indicating limited overall nucleotide similarity and highlighting the substantial divergence of the porcine adenovirus identified in this study.

To further characterise the novel porcine adenovirus, the complete genome was annotated and compared with Bat mastadenovirus WIV13 (NC_030874.1). Although the DNA polymerase of PAdV-6 clustered closely with DNA polymerase of Polar bear adenovirus 1 in phylogenetic analyses (Figure 4), this virus was not included in the comparative genomic analyses because only a partial genome sequence is currently available, precluding robust whole-genome comparisons. PAdV-6 possessed the characteristic genomic organisation of mastadenoviruses, with all major adenoviral open reading frames identified, including genes encoding structural and replication proteins. Despite this conserved genomic architecture, the PAdV-6 genome was shorter than Bat mastadenovirus WIV13, comprising 27,685 nucleotides compared with 29,162 nucleotides.

The only additional porcine virus detected in any of the samples was Porcine kobuvirus (PZ721985), which was identified in one of the piglet tissue libraries at relatively low abundance (126 RPM) (Figure 3). The virus shared 99% amino acid identity with its closest reference sequence (AXB26734.1), previously detected in rectal swabs from pigs (*Sus scrofa domesticus*) in Vietnam (Figure 5). Given the high level of sequence identity to previously described strains, this virus is not considered novel. Its low abundance and limited detection across libraries suggest it is unlikely to be the primary agent associated with the observed pathology, however, a potential contributory role in co-infection or background enteric viral presence cannot be excluded.

**Figure 5.**
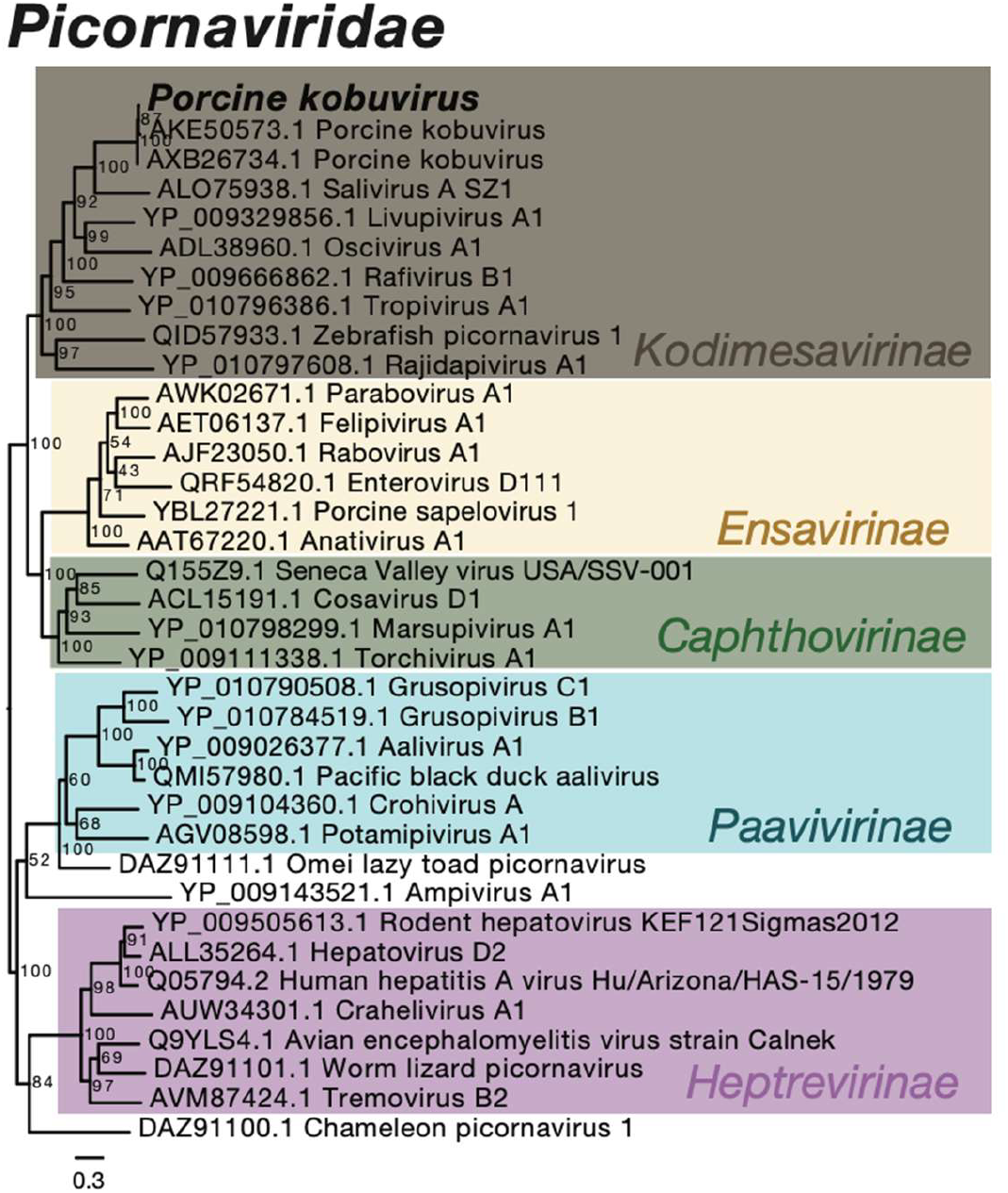
*Picornaviridae* phylogenetic tree. A maximum likelihood phylogenetic tree was constructed from representative picornavirus polyprotein transcripts. Branch lengths are proportional to the number of amino acid substitutions per site, and the tree is midpoint rooted. Ultrafast bootstrap values are noted and subfamilies are coloured.

## Discussion

Here we report the detection and genomic characterisation of a novel porcine adenovirus associated with a disease outbreak in a 320-sow high-health, pig herd in New Zealand. The outbreak was characterised by anorexia, lethargy, marked diffuse cutaneous erythema and death in neonatal piglets, together with an increased occurrence of mummified fetuses. Although PAdV-3 and PAdV-5 have previously been detected in pig faecal samples in New Zealand, with PAdV-5 also identified in river water and shellfish as a marker of porcine faecal contamination^17^ and adenovirus inclusion bodies have been an incidental histological finding in piglets^16^, no porcine adenovirus-associated disease outbreaks have previously been reported in New Zealand. The novel virus identified in this study, provisionally named PAdV-6, was highly abundant in piglet tissues and was also detected in two of the four fetal metatranscriptomic libraries. Histopathological findings were consistent with a systemic viral infection, including widespread intranuclear inclusion bodies within endothelial cells. These findings suggest active viral replication in affected tissues.

Porcine adenoviruses are generally considered low-pathogenicity viruses and have been detected in both clinically healthy and diseased pigs, where they are most commonly associated with mild, self-limiting enteric disease ^13,14^. Although porcine adenoviruses have occasionally been reported in association with reproductive failure and abortion, evidence supporting a direct causal role remains limited^41,42^. The systemic distribution of PAdV-6 together with the associated histopathological lesions, marked diffuse cutaneous erythema, increase in the rate of mummies and lack of diarrhoea, therefore represents an unusual presentation.

An important factor likely contributing to the severity of disease observed in the present outbreak is the immunological status of the affected herd. The pigs originated from a hysterectomy-derived herd established to exclude specific pathogens, resulting in animals with limited prior exposure to infectious agents. It is therefore plausible that introduction of PAdV-6 into this immunologically naïve population resulted in systemic infection and reproductive disease that would not typically occur in conventionally reared pigs with pre-existing immunity. This hypothesis is consistent with the transient nature of the outbreak, which resolved within six weeks, suggesting that herd immunity developed following viral introduction.

This interpretation is supported by previous disease events in the same herd that highlight the vulnerability of high health herds when biosecurity breaches occur. In 2021, an outbreak characterised by abortions, stillbirths, mummified fetuses, increased piglet mortality and respiratory disease was attributed to porcine cytomegalovirus following exclusion of exotic and endemic pathogens^43^. As with the present outbreak, disease severity was thought to reflect the herd’s immunologically naïve status, with losses declining as herd immunity developed^43^. The herd also experienced significant and atypical disease outbreaks associated with *Glaesseralla parasuis* servoar 5 (2021) and *Pasteurella multocida* (2022) before endemic stability was reached (Bruce Welch.,pers comm). Comparable observations to those seen in this case have also been reported in hysterectomy-derived, colostrum-deprived pigs, where a disseminated adenovirus-like infection caused severe systemic disease characterised by skin cyanosis and subcutaneous oedema in the submandibular, thoracic and abdominal regions^23^. Based on these findings, the authors suggested that the adenovirus-like agent may have been capable of transplacental transmission^23^. Together, these observations suggest that host immune status may be a critical determinant of disease severity following adenovirus infection in pigs.

Notably, only one additional porcine virus, porcine kobuvirus, was detected in the small intestine and colon of the infected piglet and only at low abundance. Porcine kobuvirus is widely distributed in pig populations and is most commonly associated with enteric infections or detected in clinically healthy animals^44^. Its low abundance and established tissue tropism make it unlikely to account for the widespread systemic pathology observed in this outbreak, although the possibility of co-infection or synergistic interactions cannot fully be excluded. In contrast, the high abundance of PAdV-6 in affected tissues together with compatible histopathological lesions, evidence of systemic infection and detection in fetal tissues strongly support its involvement in the disease process.

The evolutionary origin of PAdV-6 remains uncertain. One possibility is that it represents a previously undescribed lineage of porcine adenovirus. Porcine adenoviruses remain sparsely sampled globally ^13,14^, with relatively few complete genomes available and substantial gaps in surveillance likely to underestimate their true genetic diversity. Consequently, the apparent divergence of PAdV-6 may reflect limited sampling rather than emergence from another host species. Therefore, it is plausible that conventional health status herds in New Zealand and globally have encountered PAdV-6 without experiencing clinical disease as cross-protection has been afforded by background immunity against other porcine adenoviruses.

However, phylogenetic analysis demonstrated that PAdV-6 shared limited similarity with previously described porcine adenoviruses and instead clustered more closely with adenoviruses detected in bats and other mammals, including a highly divergent adenovirus identified in a four-month-old polar bear (*Ursus maritimus*) in Germany^38^. The polar bear cub presented with severe abdominal pain and mild diarrhoea before dying, with postmortem molecular analyses revealing systemic infection involving blood, liver, kidneys, lymph node and intestine^38^. Notably, no other polar bears in the facility tested positive for Polar bear adenovirus 1, suggesting that the infection either originated from an external reservoir or represented an opportunistic infection rather than sustained in-captive transmission^38^. Although the ecology and host range of these divergent mammalian adenoviruses remain poorly understood, these observations demonstrate that highly divergent adenoviruses can be associated with severe systemic disease following apparent cross-species transmission.

The phylogenetic placement of PAdV-6 raises the possibility that the virus may represent a relatively recent host-switching event into pigs from an as-yet unsampled mammalian reservoir. Small amounts of frozen semen from another herd have been used in this herd since creation in 2020, suggesting one potential route of virus introduction. However, any such semen stocks from the period in which the sows involved in this outbreak had been mated, had been exhausted at the time of this investigation, preventing retrospective virological screening and limiting the ability to determine whether semen contributed to virus introduction.

Alternatively, the virus may have originated from mammals present around the farm, including cattle, rodents, rabbits, hares or feral cats. Environmental contamination also represents a plausible pathway, as pigs were bedded on untreated straw that may have been contaminated with faeces or other biological material from wild mammals, including rodents, rabbits and deer, before use. Although these potential introduction routes remain speculative, they highlight the complexity of reconstructing transmission pathways once an outbreak has occurred. They also emphasise the need for broader genomic surveillance of adenoviruses in domestic swine and sympatric mammalian species to better characterise viral diversity, identify potential reservoir hosts and improve our understanding of cross-species transmission.

Overall, this study expands the known diversity of porcine adenoviruses by describing a highly divergent lineage associated with systemic infection in pigs. The application of metatranscriptomic sequencing enabled recovery of the complete viral genome while simultaneously demonstrating high viral abundance across affected tissues, providing strong evidence linking PAdV-6 with this disease outbreak. More broadly, these findings suggest that adenoviruses may represent an under-recognised cause of reproductive disease in swine, particularly in immunologically naïve populations. Although the pathogenic potential of PAdV-6 in conventional swine populations remains unknown based on the available evidence, PAdV-6 is unlikely to cause severe disease in conventional health status herds with pre-existing immunity, highlighting the importance of host immune status in determining disease outcome. Future surveillance combined with experimental infection studies will be required to determine the pathogenic potential, host range and epidemiological significance of PAdV-6 in New Zealand and internationally.

## Resource availability

Viral sequences have been submitted to GenBank under the accession numbers PZ721985-PZ721986 while raw sequencing reads are available under BioProject PRJNA1497544.

## Acknowledgements

Thanks to Rob Fairley (Awanui Veterinary, Pathologist) for reviewing the histology, Mark Bestbier (Animal Health Laboratory, Pathologist) for technical support providing fetus tissue samples and managing subcontracted tests. Also thanks to Jason Smith and Keanan Sylvester (AHL technicians) for the described PCR testing of FFPE tissues and fresh tissue samples for viruses and Andrew Wilson (AHL Genomics Manager) for providing initial sequence data from PCR amplicons.

## Supplementary information

**Supplementary Figure 1. PCR screening of fetal and neonatal porcine tissues for adenovirus-like and related viral DNA**. (a) Formalin-fixed paraffin-embedded (FFPE) tissues from a symptomatic, deceased neonatal piglet showing intranuclear inclusion bodies. DNA was extracted from 10 µm paraffin scrolls derived from spleen and lymph node (lanes 1, 5), heart, lung, liver, and kidney (lanes 2, 6), and small intestine and colon (lanes 3, 7). Avian adenovirus DNA and equine herpesvirus type 1 DNA were included as positive controls (lanes 4 and 8, respectively). (b) Pan-adenovirus nested PCR screening of nucleic acids extracted from homogenised tissues of mummified fetuses. Tissues include lung (lanes 1, 4, 8, 11), kidney (lanes 2, 6, 13), liver (lanes 9, 12), and brain (lanes 3, 7, 10, 14), corresponding to fetus 1 (lanes 1–3), fetus 2 (lanes 4–7), fetus 3 (lanes 8–10), and fetus 4 (lanes 11–14). Avian adenovirus DNA was used as a positive control (P).(c) PCR amplification of thymus (lanes 1, 3, 5, 7) and spleen (lanes 2, 4, 6, 8) tissue extracts from mummified fetuses collected during the outbreak period, including fetus 1 (lanes 1–2), fetus 2 (lanes 3–4), fetus 3 (lanes 5–6), and fetus 4 (lanes 7–8), using novel nPAd-long primers. All PCR products were resolved on 1.5% TAE agarose gels. ne, negative extraction control; nt, negative template control; M, 100 bp molecular weight marker (ThermoFisher Scientific).

**Supplementary Table 1. Information around the tissue samples that were collected and analysis that was done**.

**Supplementary Table 2. PCR assays used for targeted screening of viral pathogens**.

**Supplementary Table 3. Molecular diagnostic screening of fetal tissues by PCR for DNA and RNA viruses associated with porcine prenatal deaths and mummification**.

**Supplementary Table 4. Screening prenatal fetal tissues for adenoviruses using PCR methods**.

**Supplementary Table 5. Predicted open reading frames (ORFs) encoded by PAdV-6 and their amino acid similarity to mammalian adenoviruses**.

